# Characterization of nuclear mitochondrial insertions in canine genome assemblies

**DOI:** 10.1101/2024.09.13.612826

**Authors:** Peter Z. Schall, Jennifer R.S. Meadows, Fabian Ramos-Almodovar, Jeffrey M. Kidd

**Affiliations:** Department of Human Genetics, University of Michigan, Ann Arbor, Michigan, USA. 48109; Department of Medical Biochemistry and Microbiology, Uppsala University, Uppsala, Sweden. 75132; SciLifeLab, Uppsala University, Uppsala, Sweden. 75132; Department of Computational Medicine & Bioinformatics, University of Michigan, Ann Arbor, Michigan, USA. 48109

**Keywords:** Canine, nuclear mitochondria insertion, Numt

## Abstract

**Background:** The presence of mitochondrial sequences in the nuclear genome (Numts) confounds analyses of mitochondrial sequence variation and is a potential source of false positives in disease studies. To improve the analysis of mitochondrial variation in canines, we completed a systematic assessment of Numt content across genome assemblies, canine populations and the carnivore lineage.

**Results:** Centering our analysis on the UU_Cfam_GSD_1.0/canFam4/Mischka assembly, a commonly used reference in dog genetic variation studies, we find a total of 321 Numts, located throughout the nuclear genome and encompassing the entire sequence of the mitochondria. Comparison to 14 canine genome assemblies identified 63 Numts with presence-absence dimorphism among dogs, wolves, and a coyote. Further, a subset of Numts were maintained across carnivore evolutionary time (arctic fox, polar bear, cat), with 8 sequences likely more than 10 million years old, and shared with the domestic cat. On a population level, using structural variant data from the Dog10K Consortium for 1,879 dogs and wolves, we identified 11 Numts that are absent in at least one sample as well as 53 Numts that are absent from the Mischka assembly.

**Conclusions:** We highlight scenarios where the presence of Numts is a potentially confounding factor and provide an annotation of these sequences in canine genome assemblies. This resource will aid the identification and interpretation of polymorphisms in both somatic and germline mitochondrial studies in canines.

## Introduction

Since its origin as an organelle, components of the mitochondrial genome (mtDNA) have been repeatedly transferred to the eukaryotic nuclear genome [1]. Nuclear mitochondrial (Numt) sequences are segments of the nuclear genome with a recognizable origin from the mitochondrial genome [2]. Hybridization experiments from 40 years ago suggested the presence of mitochondrial-like sequences in the nuclear genome and the advent of large-scale genome sequencing confirmed the ubiquitous presence of Numts across eukaryotes [3]. Analysis of diverse species indicate a wide range in the number and cumulative length of Numts across taxa, with some species showing evidence of post-insertion Numt amplification [2,4-9]. Numts can be useful markers for evolutionary studies and may show presence-absence dimorphism within a species [10-14]. Recent large-scale analyses in humans confirm that Numt formation is an ongoing process with new Numt insertions found in normal somatic tissues and tumors as well as being transmitted through the germline [15,16]. Analysis of rare Numts in humans indicates that Numt breakpoints are enriched in non-coding segments of the mitochondria genome, Numt insertions are depleted near genes in the nuclear genome, and Numt formation involves multiple molecular mechanisms related to genome stability, DNA damage, and repair [15].

The presence of Numts in the nuclear genome complicates the analysis of mitochondrial sequence variation. Sequence differences between Numts and the mitochondria can be misinterpreted as mitochondrial mutations [17,18] or heteroplasmies [19-22]. Numts may also be associated with errors in genome assembly, resulting from false contig joins between nuclear and mitochondrial sequences that lead to the artifactual presence of large, mitochondrially derived sequences in an assembled nuclear chromosome [23-25].

Domestic dogs (*Canis lupus familiaris*) are a valuable model for studies of evolution and disease [26,27]. Similar to other mammals, dog genetic diseases caused by mutations in the nuclear genome are more widely studied than mitochondrial disease. However, multiple disease-causing mutations in dog mtDNA and affecting mitochondrial function have been described [28]. In canines, mitochondrial changes have also been associated with tumor progression [29,30] and the evolution of the clonally-inherited canine transmissible venereal tumor (CTVT) [31-33]. Failure to account for Numts in the canine genome may confound the interpretation of canine mitochondrial variation.

Analysis of nuclear DNA from sperm heads [34], as well as bioinformatic analysis of the initial canine genome assembly [35,36], confirmed the presence of Numts in dogs. Recent advances in long-read genome sequencing have led to the publication of multiple canine genomes, but analysis of Numts across assemblies, including the characterization of dimorphic Numts, has been limited. As part of their assembly quality control procedure, Edwards et al. performed a systematic assessment of Numts in their basenji genome assemblies, finding patterns generally consistent with previous studies of Numts in mammals and limited dimorphism among assemblies [24]. To analyze mitochondrial variation in samples sequenced by the Dog10K Consortium, we previously identified large, high-identity Numts in the UU_Cfam_GSD_1.0/canFam4 assembly derived from a German Shepherd Dog named Mischka [37,38]. In this study, we provide a systematic analysis of Numts in 15 genomes assemblies from dogs, wolves, and a coyote and assess Numt sharing across Carnivora. We characterize multiple Numts that differ among assemblies and additionally identify dimorphic Numts using Illumina sequencing data from 1,879 individuals. These data will aid future studies of somatic and germline mitochondrial variation in canines.

## Methods

### Identification of Numts in genome assemblies

We identified nuclear mitochondrial sequences (Numts) in canine genome reference assemblies based on the procedure previously used for analysis of the human genome [39]. First, the canine mitochondrial reference genome [40] (NC_002008.4) was searched against the genome assembly using the bl2seq functionality of blastn in the NCBI BLAST+ package [41], version 2.10.0. The search was performed with a scoring function of +2 for matches, -3 for mismatches, -5 for gap opening, and -2 for gap extension. Only high scoring pairs (HSPs) with an e-value less than 0.001 were retained. HSPs with coordinates within 2,000 bp in both the genome and mitochondria sequence, with consistent orientation, were merged into Numts regions (‘assembled Numts’ in the terminology of [39]). To refine HSP length and identity relative to the circular mitochondrial genome, a final set of HSPs was identified by searching the sequence of each merged Numt segment with a query consisting of the mitochondrial reference sequence concatenated twice. Numt length was tabulated based on the number of aligned mitochondrial bp in each HSP. The Numt content of 15 genome assemblies derived from 14 individuals was assessed (Table S1), including: two basenjis [24], two Bernese Mountain dogs [42], two different assemblies created from the same boxer [43,44], a Cairn terrier [42], a dingo [45], two German Shepherd Dogs [37,46], a Great Dane [47], a Labrador retriever [42], two wolves [48,49], and a coyote [49]. Analysis of the coyote assembly was performed using the coyote mitochondrial genome sequence [50] (NC_008093.1). Statistics were tabulated from Numt HSPs and merged Numts identified in each assembly, stratified by sequences assigned to assembled nuclear chromosomes (i.e., chr1-chr38 and chrX) or to unplaced contig sequences not assigned to an assembled chromosome.

### Identification of Numt differences using multiple assemblies

Numts that differ between assemblies were identified by intersecting Numt HSPs with insertion and deletion variants found between assemblies. First, each assembly was aligned to the Mischka genome using minimap2 version 2.26 with option -x asm20 [51,52]. Insertion and deletion variants were identified using the paftools.js call command to identify variants in assembly segments covered by a single long alignment (-l 10000 and -L 50000 options). Insertions and deletions were converted to BED format and intersected with Numt HSPs annotated in each assembly using bedtools version 2.26.0 [53], reporting only variants that overlapped at least 90% of the Numt HSP (-f 0.90 option). The variant was classified as a large structural variant if the detected insertion or deletion was more than 1,000 bp longer than the Numt HSP.

### Comparison with outgroup genome assemblies

To estimate Numt age, we searched for the presence of merged Mischka Numt loci in the arctic fox [54] (*Vulpes lagopus*, GCF_018345385.1), polar bear [55] (*Ursus maritimus*, GCF_017311325.1), and cat [56](*Felis catus*, felCat9/GCF_000181335.3) genome assemblies. Each locus, along with 10 kbp of flanking sequence on each side, was extracted from the Mischka genome and searched against the polar bear and cat assemblies using BLAT [57] with options -stepSize=5 -repMatch=2253 -minScore=20 -minIdentity=0. The resulting hits were filtered to retain alignments that covered at least 2,000 bp from the left and the right flanks as well as at least 80% of the Numt. To confirm hits, candidate loci were extracted from the arctic fox, polar bear, and cat assemblies and then queried against the dog mitochondrial sequence as described above.

### Identification of polymorphic Numts in Dog10K samples

Structural variants identified and genotyped in 1,879 dogs and wolves by the Dog10K consortium were retrieved from Meadows et al. [38]. Deletion structural variants were intersected with the merged Mischka Numt loci using bedtools, requiring a reciprocal overlap of 90% (-r 0.9).

Insertion variants were extracted from the VCF file, splitting by those completely assembled and those with only flanking left and right sequences. Complete sequences were converted to fasta format, using chromosome:position as identifiers. Incomplete sequences were joined with a string of 10 N’s between the left and right sequences and outputted to fasta format. The two resultant fasta files were concatenated and analyzed using numtfinder [24] to identify non-reference Numts, using the NC_002008.4 mitochondrial sequence, with the addition of the *circle=T* flag. The number of Numts present per sample was determined based on the genotypes in the Dog10K VCF file and summarized by category including breed dogs, dogs of mixed origin or that are not recognized by any international registering body (labeled as Mixed/Other), village dogs, and wolves.

## Results

### Identification of nuclear mitochondrial sequences in the Mischka UU_Cfam_GSD_1.0/canFam4 genome reference

The UU_Cfam_GSD_1.0 assembly [37], derived from a German Shepherd Dog named Mischka and labeled as canFam4 in the UCSC Genome Browser, has emerged as the main reference used for the analysis of canine genome variation [38]. We therefore first annotated nuclear mitochondrial sequences (Numts) in the Mischka assembly following the procedure previously used for the analysis humans and other species [39]. We identified a total of 321 Numt HSPs, with five HSPs located on assembled contigs that are not integrated into the chromosomal-level assembly (Table 1, Table S2, and Table S3). This includes a full-length copy of the mitochondria genome with high sequence identity that makes up chrUn_JAAHUQ010000987v1 and is likely the result of assembly error. We identified 316 Numt HSPs encompassing 200,108 mitochondrial bp on the assembled chromosomes. These 316 segments can be merged into 243 Numt loci which are located across all 38 autosomes and the X chromosome (Figure 1). A similar distribution of Numts was found across 14 other canine genome assemblies (Table S2, Table S3, and Figure S1). The Numt segments encompass the entire mitochondrial genome, with reduced coverage found in the D-loop which has been previously found to be depleted in Numts across primates and which contains a short repeat sequence that is highly variable in canines [39,58] (Figure 2).

**Table 1.**
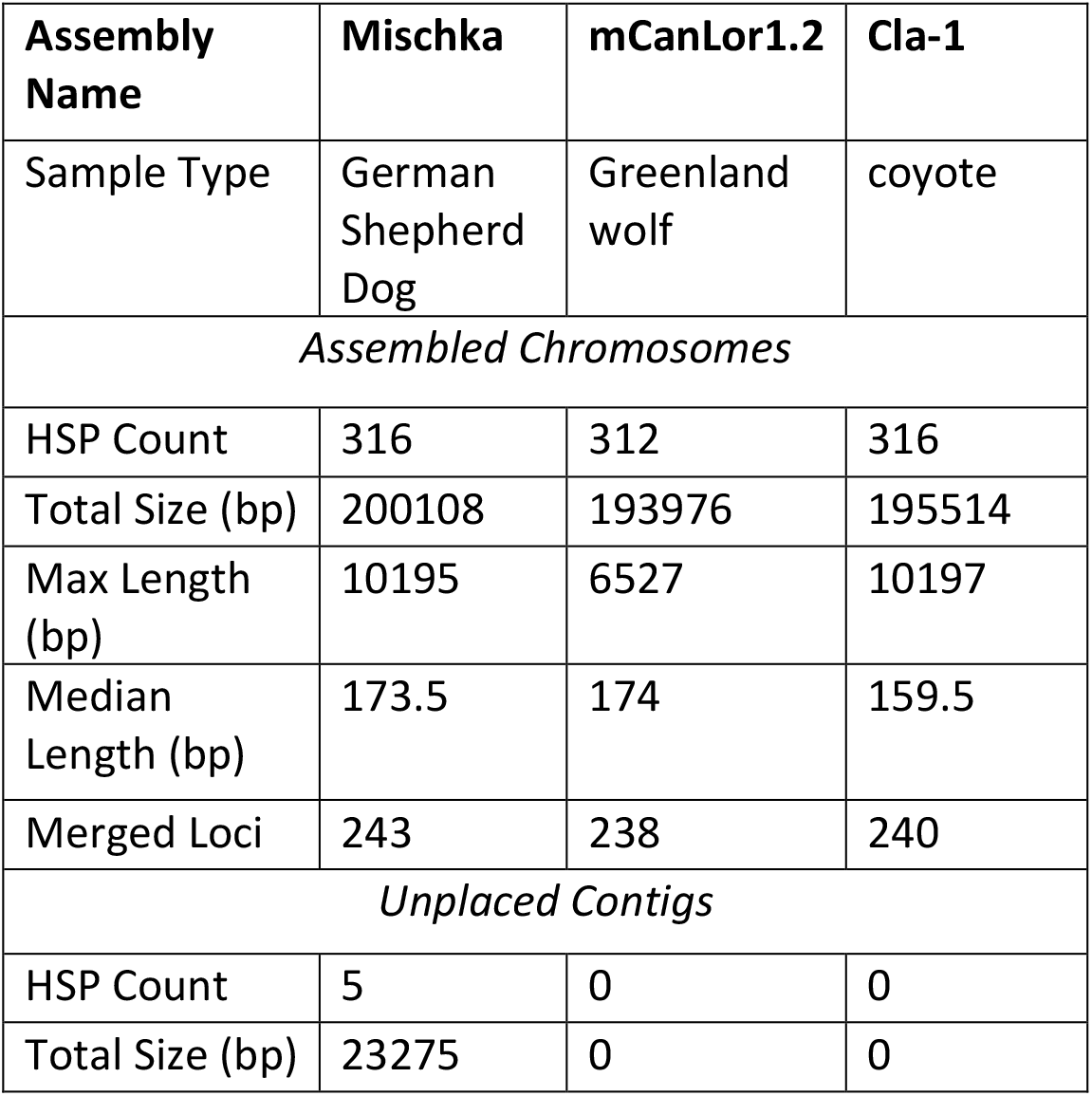
Summary of Numts identified in genome assemblies from a dog, wolf, and coyote.

| **Assembly Name** | **Mischka** | **mCanLor1.2** | **Cla-1** |
| --- | --- | --- | --- |
| Sample Type | German Shepherd Dog | Greenland wolf | coyote |
| *Assembled Chromosomes* |  |  |  |
| HSP Count | 316 | 312 | 316 |
| Total Size (bp) | 200108 | 193976 | 195514 |
| Max Length (bp) | 10195 | 6527 | 10197 |
| Median Length (bp) | 173.5 | 174 | 159.5 |
| Merged Loci | 243 | 238 | 240 |
| *Unplaced Contigs* |  |  |  |
| HSP Count | 5 | 0 | 0 |
| Total Size (bp) | 23275 | 0 | 0 |

**Figure 1.**
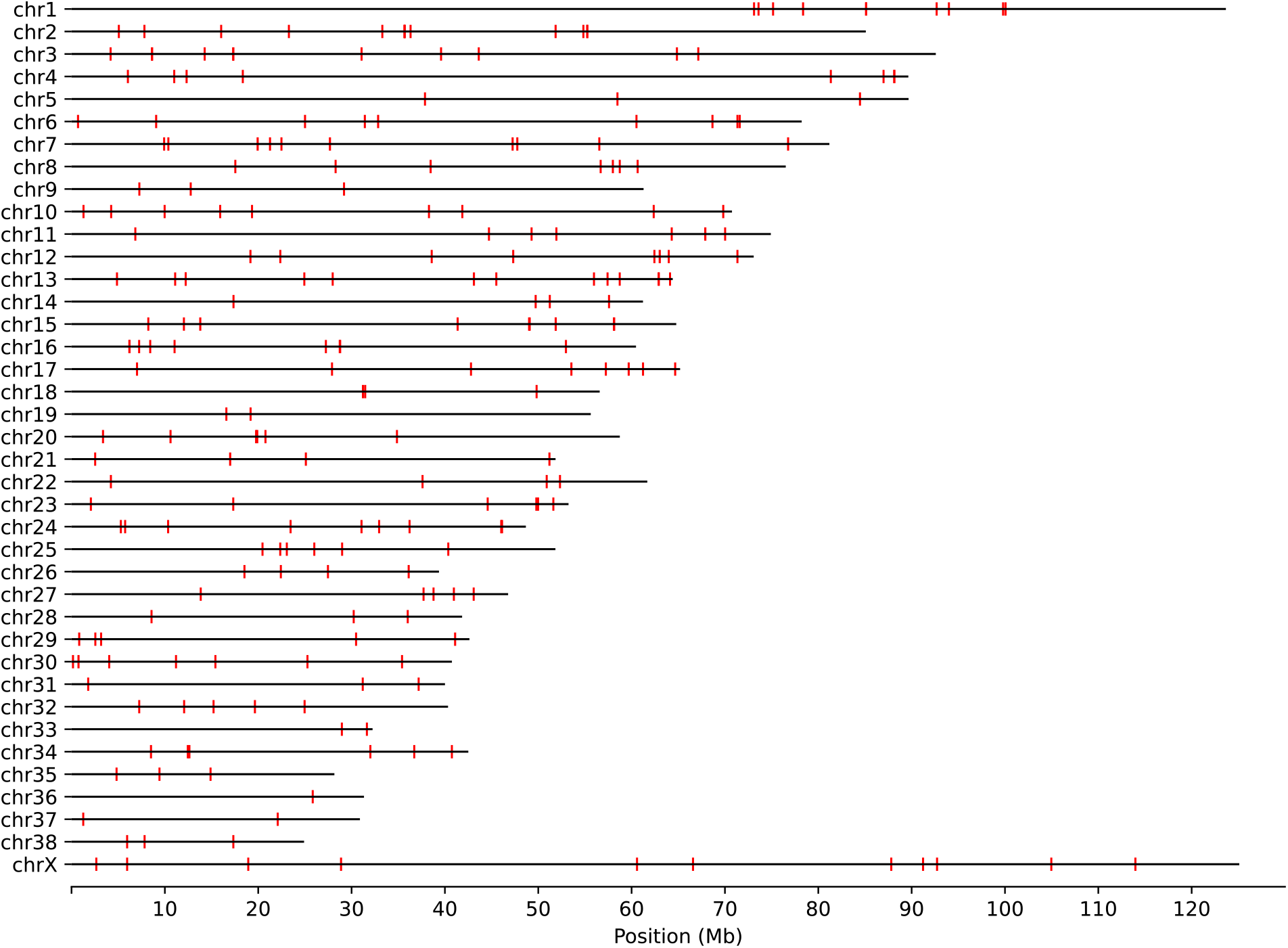
Position of Numts in the Mischka genome. The position of 243 merged Numt loci along the assembled nuclear chromosomes in the Mischka (UU_Cfam_GSD_1.0/canFam4) genome is shown. The location of each Numt is indicated by a red box.

**Figure 2.**
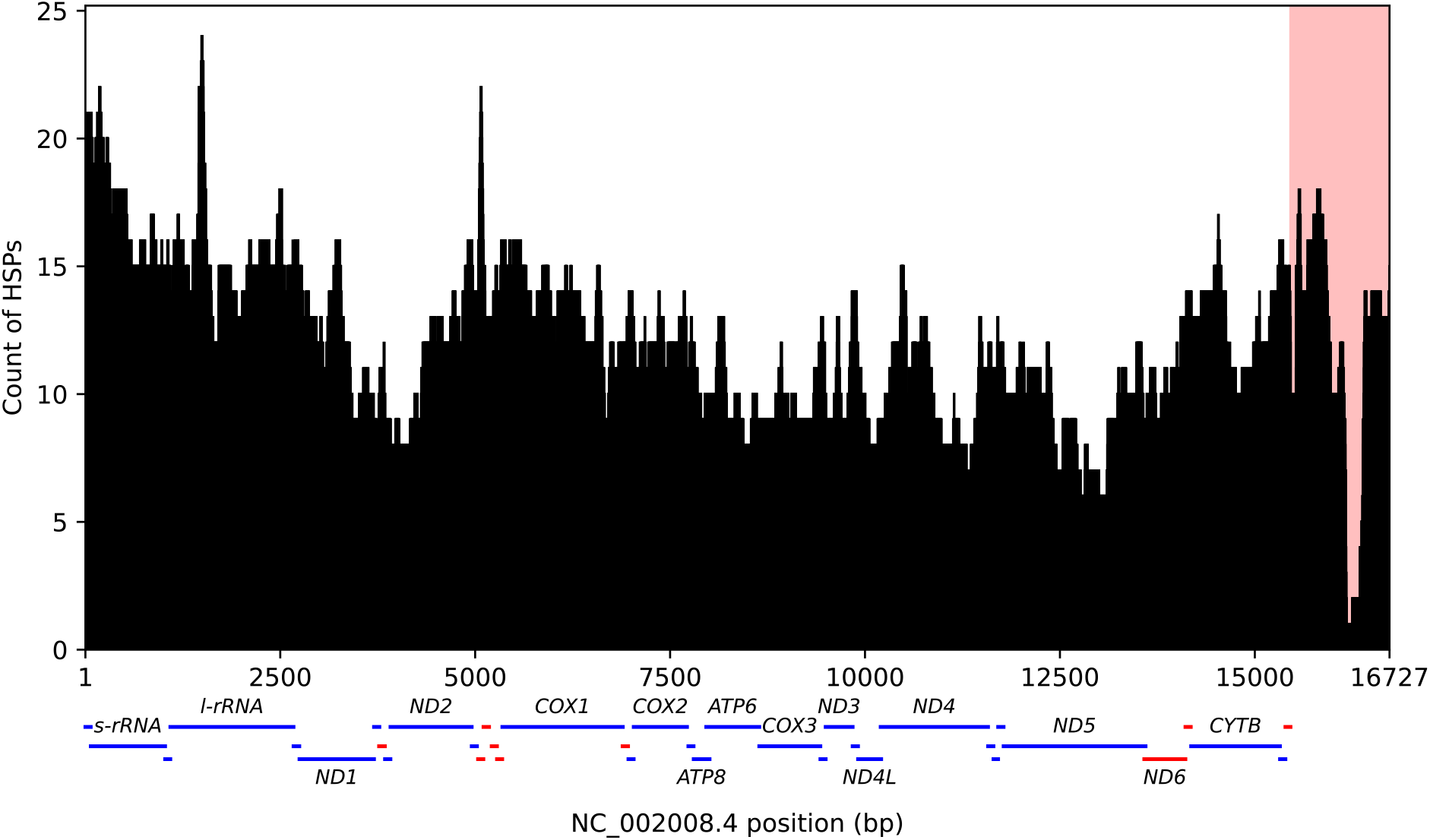
Coverage of canine mtDNA along the assembled Mischka genome. The total coverage along the mitochondria is shown for the 316 Numt HSPs identified in the assembled chromosomes from Mischka. The shaded pink region corresponds to the mitochondrial D-loop. The position of mitochondrial genes on the forward (blue) and reverse (red) strand is depicted below the figure. The names of rRNA and protein-coding genes are given.

It is not straight forward to compare the raw counts of annotated Numt HSPs between genomes. For example, the longest Numt HSP identified in the Mischka assembled chromosomes is found on chr34, encompasses 10,195 mitochondrial bp with an identity of 83.14%, and is also present in the mCanLor1.2 wolf genome. However, in mCanLor1.2 this long Numt is disrupted by a LINE-1 insertion that occurred after Numt integration (Figure S2) and is therefore reported as separate HSPs.

### Dimorphic Numts between Mischka and other genome assemblies

We searched for Numts that have presence-absence dimorphism between Mischka and other genome assemblies from dogs, wolves, and a coyote. First, we identified a total of 32 Numt HSPs (14,528 total bp) annotated in Mischka that appear to be absent in another assembly (Table S4). This includes 9 Numt HSPs that overlap with larger structural variants identified between assemblies and 23 Numts that overlap with deletion variants with a length within 1 kbp of the Numt size. Of the 23 cleanly dimorphic Numts, 6 were absent only in the coyote. One Numt, a 195 bp HSP with 98.5% identity, was absent in every other assembly analyzed. As expected, the dimorphic Numts have a higher identity to the reference mitochondria than those which are fixed in all samples (p < 0.0001, Figure 3). Considering more distantly related members of the carnivore linage, 169 of the 243 merged Numts present on assembled chromosomes in Mischka are found in the arctic fox (*Vulpes lagopus*) genome while 20 are present in the polar bear (*Ursus maritimus)* genome and 8 are present in the cat (*Felis catus)* genome, indicating that a subset of Numts have an ancient origin dating to the initial diversification of the order Carnivora (Table S5).

**Figure 3.**
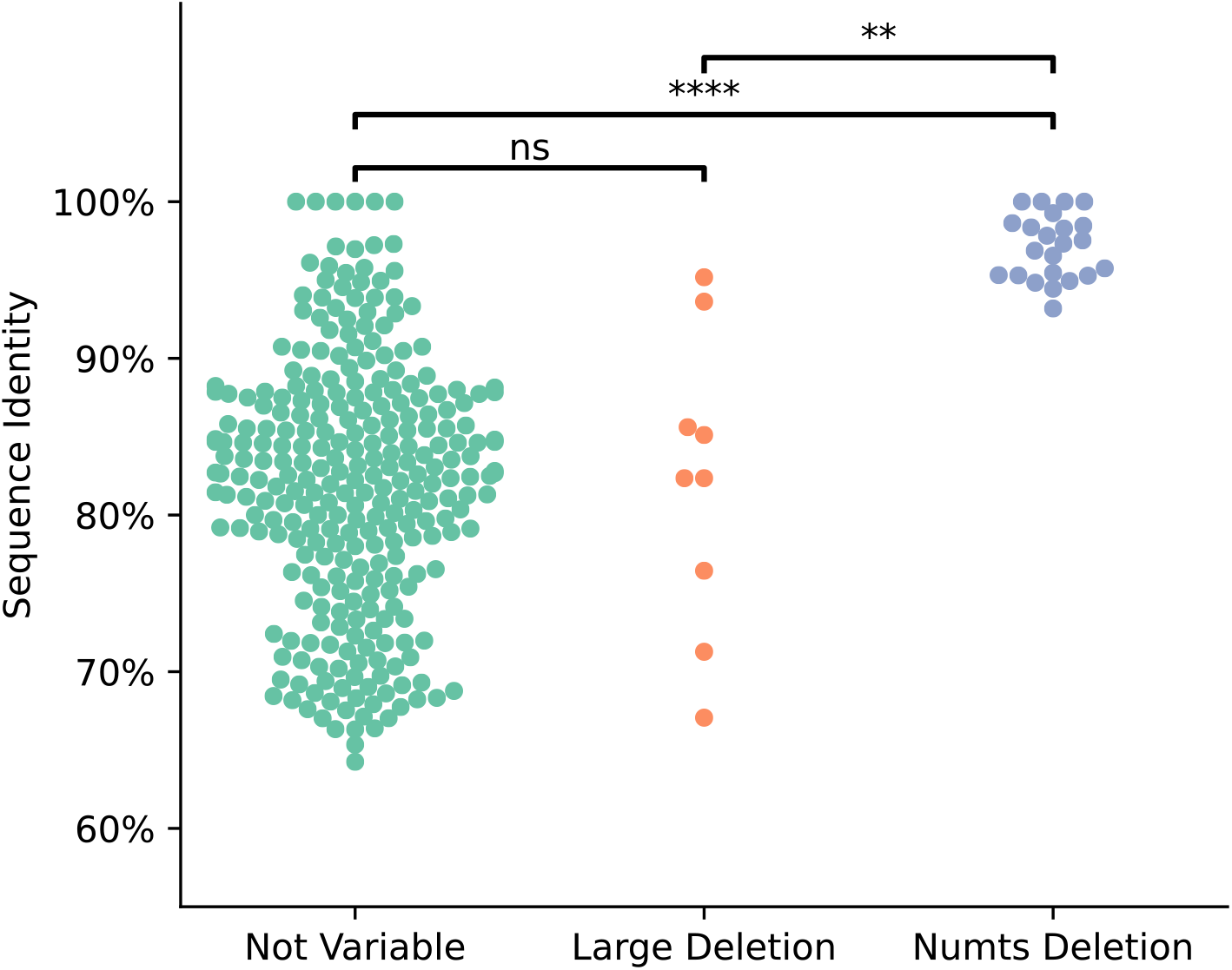
Dimorphic Numts have a higher mitochondrial sequence identity than fixed Numts. A swam plot of sequence identity relative to the mitochondrial reference genome is shown for Numts annotated in the Mischka assembly, values are plotted for 284 Numts that are found in each assembly, 9 Numts that overlap larger deletions, and 23 Numts that are absent in other assemblies. Sequence identities were compared across categories using Welch’s unequal variances t-test: n.s: not significant, ** p < 0.01, **** p < 0.0001.

We additionally searched for Numts that appear as insertions in the other assemblies relative to Mischka and identified 31 loci (Table S6). Six of these correspond to large structural variants that include a Numt sequence, while 25 are insertion differences corresponding to Numts. These 25 loci include five Numts that were found only in the coyote, as well as a 3,280 bp segment present only in Sandy, the dingo. Together, analysis of these assemblies identifies 63 Numts that show presence-absence dimorphism among dogs, wolves, and coyotes.

### Comparison with 1,879 samples analyzed by the Dog10K consortium

The Dog10K consortium identified structural variants in 1,879 diverse canines based on alignment of Illumina sequencing data to the Mischka genome assembly [38]. The Dog10K deletions include 11 variants that have a reciprocal 90% overlap with a merged Numt locus annotated in Mischka (Table S7). This includes one Numt not identified as variable based on the analysis of 14 genome assemblies described above.

The Dog10K structural variant collection also includes insertions along with assemblies of the insertion sequence. Since the Dog10K data is derived from Illumina reads, the full sequence of large insertions (⪆ 200 bp) could not be resolved and is represented as partial segments extending into the variant from each edge. We compared the reported sequence of each insertion with the mitochondrial reference and identified 53 insertions that correspond to Numts (Table S8). This includes 7 variants where the insertion sequence is only partially assembled. Of the 53 insertions, 15 were also identified in our analysis of the 14 additional genome assemblies. Dog10K samples contained a median of 4 Numts, with a range of 0 to 11 (Figure 4). Of the 1,879 samples, only 12 did not have any of the 53 insertions. Delineating by canine category, Breed Dogs (n=1,575) displayed a median of 4 insertions per sample (range=0-11), Village Dogs (n=237) median of 5 (range=1-11), and Wolves (n=55) with a median of 7 (range=3-11).

**Figure 4.**
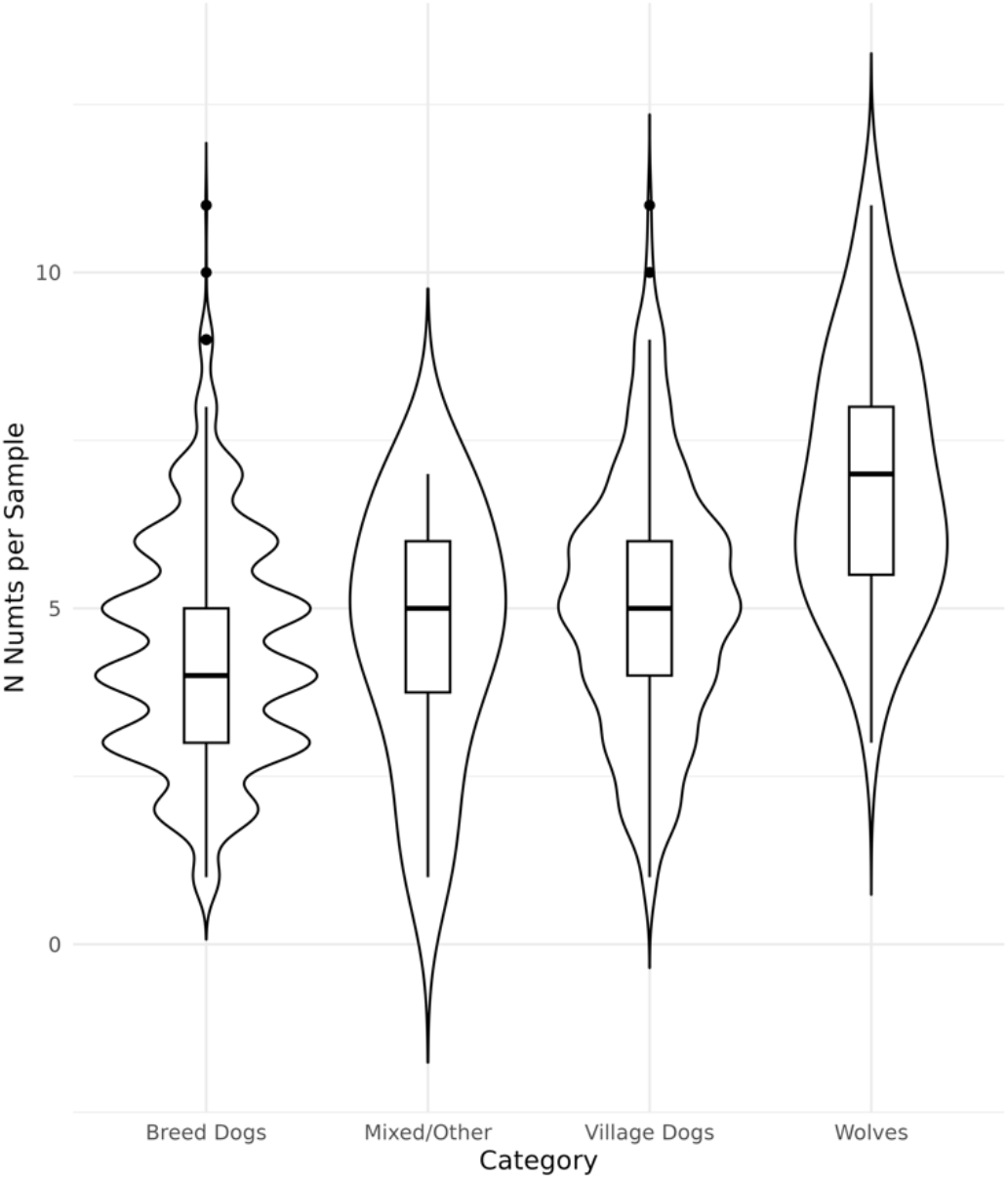
Numts insertions identified in Dog10K samples. Insertions identified in 1,879 canines by the Dog10K consortium were analyzed to identify 53 insertions that correspond to Numts. Violin plots and boxplots are shown of the number of Numts present per sample, stratified by category: Breed Dogs (n=1,575), Mixed/Other (n=12) Village Dogs (n=237) and Wolves (n=55).

## Discussion

The Mischka genome [37] (UU_Cfam_GSD_1.0/canFam4), derived from a German Shepherd Dog, has emerged as a common reference genome used for studies of canine variation, making annotation of Numts in this genome a valuable resource for the canine genomics community [38]. We identified 321 Numt HSPs in the Mischka genome that in total encompass the entire mitochondrial genome sequence. This includes a full-length representation of the mitochondria genome present on an unplaced sequence that is included in the Mischka assembly but is not localized to an assembled chromosome.

Knowing the location and types of Numts within the analyzed reference sequence is key to the success of future research projects. From a nuclear genome perspective, a Numt black list can highlight those regions likely to negatively impact short read mapping or cell-free mtDNA detection [3,35,59,60]. From the mitogenome perspective, if not accounted for, the full-length Numt spanning an unplaced contig in the Mischka genome will act as a decoy and siphon reads from the true mitochondrial genome. This has the potential to lead to underestimates of mitochondrial copy number and to reduce the measurable allele fraction of true mitochondria variants. Additionally, false levels of mitochondrial heteroplasmy will be estimated if Numt sequences are not accounted for in mitochondrial mapping and variant calling pipelines. This problem can be address through the incorporation of a Numt reference list to mitochondrial mapping software. To facilitate such analyses, we have collated the Numt locations in the Mischka, Tasha, canFam3.1, Zoey and mCanLor1.2 genomes into an UCSC Genome Browser Tack Hub [61,62].

To put our annotation of Mischka into a larger context, we identified Numts in 14 other genome assemblies. Previous analysis showed that canine genome assemblies differ markedly in the representation of duplicated sequences on unplaced chromosome contigs [63]. The same is true for Numts: three assemblies show substantial Numt representation on unlocalized contigs while the others contain little or no Numt sequence on unplaced contigs. These discrepancies likely reflect differences in the assembly algorithms and filtering strategies employed for each genome.

Of the Numts annotated in Mischka, 32 were absent in at least one other canine genome assembly. Since genomic DNA is generally not available from the samples used to construct each assembly, we were not able to experimentally validate the annotated Numts. However, we note that the dimorphic Numts have a higher sequence identity with the reference canine mitochondria genome, consistent with a more recent origin. Analysis of a diverse collection of 1,879 canines additionally identified 11 Numts loci that are deleted in at least one sample, as well as 53 insertions that correspond to non-reference Numts. Together, these observations indicate that some Numts formed recently in canine evolution and have not become fixed in canines. The presence of polymorphic Numts is a confounding factor that must be accounted for in studies of mitochondrial sequence variation, mitochondrial heteroplasmy, and somatic mutation in canines.

## Data availability

UCSC Track Hub of Numts https://github.com/KiddLab/dog-numts-tracks

HSP identification and merging script: https://github.com/jmkidd/refNUMTS

## Author Contributions

Conceptualization: P.Z.S., J.R.S.M., and J.M.K, Methodology: P.Z.S., F.R.A., and J.M.K., Analysis: P.Z.S., J.R.S.M., F.R.A., and J.M.K., Manuscript writing: P.Z.S., J.R.S.M., and J.M.K.

## References

1. Roger, A.J.; Munoz-Gomez, S.A.; Kamikawa, R. The Origin and Diversification of Mitochondria. Curr Biol 2017, 27, R1177–R1192, doi:10.1016/j.cub.2017.09.015.

2. Lopez, J.V.; Yuhki, N.; Masuda, R.; Modi, W.; O’Brien, S.J. Numt, a recent transfer and tandem amplification of mitochondrial DNA to the nuclear genome of the domestic cat. J Mol Evol 1994, 39, 174–190, doi:10.1007/BF00163806.

3. Xue, L.; Moreira, J.D.; Smith, K.K.; Fetterman, J.L. The Mighty NUMT: Mitochondrial DNA Flexing Its Code in the Nuclear Genome. Biomolecules 2023, 13, doi:10.3390/biom13050753.

4. Pamilo, P.; Viljakainen, L.; Vihavainen, A. Exceptionally high density of NUMTs in the honeybee genome. Mol Biol Evol 2007, 24, 1340–1346, doi:10.1093/molbev/msm055.

5. Hebert, P.D.N.; Bock, D.G.; Prosser, S.W.J. Interrogating 1000 insect genomes for NUMTs: A risk assessment for estimates of species richness. PLoS One 2023, 18, e0286620, doi:10.1371/journal.pone.0286620.

6. Blanchard, J.L.; Schmidt, G.W. Pervasive migration of organellar DNA to the nucleus in plants. J Mol Evol 1995, 41, 397–406, doi:10.1007/BF00160310.

7. Stupar, R.M.; Lilly, J.W.; Town, C.D.; Cheng, Z.; Kaul, S.; Buell, C.R.; Jiang, J. Complex mtDNA constitutes an approximate 620-kb insertion on Arabidopsis thaliana chromosome 2: implication of potential sequencing errors caused by large-unit repeats. Proc Natl Acad Sci U S A 2001, 98, 5099–5103, doi:10.1073/pnas.091110398.

8. Sacerdot, C.; Casaregola, S.; Lafontaine, I.; Tekaia, F.; Dujon, B.; Ozier-Kalogeropoulos, O. Promiscuous DNA in the nuclear genomes of hemiascomycetous yeasts. FEMS Yeast Res 2008, 8, 846–857, doi:10.1111/j.1567-1364.2008.00409.x.

9. Kim, J.H.; Antunes, A.; Luo, S.J.; Menninger, J.; Nash, W.G.; O’Brien, S.J.; Johnson, W.E. Evolutionary analysis of a large mtDNA translocation (numt) into the nuclear genome of the Panthera genus species. Gene 2006, 366, 292–302, doi:10.1016/j.gene.2005.08.023.

10. Thomas, R.; Zischler, H.; Paabo, S.; Stoneking, M. Novel mitochondrial DNA insertion polymorphism and its usefulness for human population studies. Hum Biol 1996, 68, 847–854.

11. Lang, M.; Sazzini, M.; Calabrese, F.M.; Simone, D.; Boattini, A.; Romeo, G.; Luiselli, D.; Attimonelli, M.; Gasparre, G. Polymorphic NumtS trace human population relationships. Hum Genet 2012, 131, 757–771, doi:10.1007/s00439-011-1125-3.

12. Dayama, G.; Emery, S.B.; Kidd, J.M.; Mills, R.E. The genomic landscape of polymorphic human nuclear mitochondrial insertions. Nucleic Acids Res 2014, 42, 12640–12649, doi:10.1093/nar/gku1038.

13. Dayama, G.; Zhou, W.; Prado-Martinez, J.; Marques-Bonet, T.; Mills, R.E. Characterization of nuclear mitochondrial insertions in the whole genomes of primates. NAR Genom Bioinform 2020, 2, lqaa089, doi:10.1093/nargab/lqaa089.

14. Uvizl, M.; Puechmaille, S.J.; Power, S.; Pippel, M.; Carthy, S.; Haerty, W.; Myers, E.W.; Teeling, E.C.; Huang, Z. Comparative Genome Microsynteny Illuminates the Fast Evolution of Nuclear Mitochondrial Segments (NUMTs) in Mammals. Mol Biol Evol 2024, 41, doi:10.1093/molbev/msad278.

15. Wei, W.; Schon, K.R.; Elgar, G.; Orioli, A.; Tanguy, M.; Giess, A.; Tischkowitz, M.; Caulfield, M.J.; Chinnery, P.F. Nuclear-embedded mitochondrial DNA sequences in 66,083 human genomes. Nature 2022, 611, 105–114, doi:10.1038/s41586-022-05288-7.

16. Zhou, W.; Karan, K.R.; Gu, W.; Klein, H.U.; Sturm, G.; De Jager, P.L.; Bennett, D.A.; Hirano, M.; Picard, M.; Mills, R.E. Somatic nuclear mitochondrial DNA insertions are prevalent in the human brain and accumulate over time in fibroblasts. PLoS Biol 2024, 22, e3002723, doi:10.1371/journal.pbio.3002723.

17. Yao, Y.G.; Kong, Q.P.; Salas, A.; Bandelt, H.J. Pseudomitochondrial genome haunts disease studies. J Med Genet 2008, 45, 769–772, doi:10.1136/jmg.2008.059782.

18. Wallace, D.C.; Stugard, C.; Murdock, D.; Schurr, T.; Brown, M.D. Ancient mtDNA sequences in the human nuclear genome: a potential source of errors in identifying pathogenic mutations. Proc Natl Acad Sci U S A 1997, 94, 14900–14905, doi:10.1073/pnas.94.26.14900.

19. Maude, H.; Davidson, M.; Charitakis, N.; Diaz, L.; Bowers, W.H.T.; Gradovich, E.; Andrew, T.; Huntley, D. NUMT Confounding Biases Mitochondrial Heteroplasmy Calls in Favor of the Reference Allele. Front Cell Dev Biol 2019, 7, 201, doi:10.3389/fcell.2019.00201.

20. Wei, W.; Pagnamenta, A.T.; Gleadall, N.; Sanchis-Juan, A.; Stephens, J.; Broxholme, J.; Tuna, S.; Odhams, C.A.; Genomics England Research, C.; BioResource, N.; et al. Nuclear-mitochondrial DNA segments resemble paternally inherited mitochondrial DNA in humans. Nat Commun 2020, 11, 1740, doi:10.1038/s41467-020-15336-3.

21. Lutz-Bonengel, S.; Niederstatter, H.; Naue, J.; Koziel, R.; Yang, F.; Sanger, T.; Huber, G.; Berger, C.; Pflugradt, R.; Strobl, C.; et al. Evidence for multi-copy Mega-NUMTs in the human genome. Nucleic Acids Res 2021, 49, 1517–1531, doi:10.1093/nar/gkaa1271.

22. Laricchia, K.M.; Lake, N.J.; Watts, N.A.; Shand, M.; Haessly, A.; Gauthier, L.; Benjamin, D.; Banks, E.; Soto, J.; Garimella, K.; et al. Mitochondrial DNA variation across 56,434 individuals in gnomAD. Genome Res 2022, 32, 569–582, doi:10.1101/gr.276013.121.

23. Shi, H.; Xing, Y.; Mao, X. The little brown bat nuclear genome contains an entire mitochondrial genome: Real or artifact? Gene 2017, 629, 64–67, doi:10.1016/j.gene.2017.07.065.

24. Edwards, R.J.; Field, M.A.; Ferguson, J.M.; Dudchenko, O.; Keilwagen, J.; Rosen, B.D.; Johnson, G.S.; Rice, E.S.; Hillier, D.; Hammond, J.M.; et al. Chromosome-length genome assembly and structural variations of the primal Basenji dog (Canis lupus familiaris) genome. BMC Genomics 2021, 22, 188, doi:10.1186/s12864-021-07493-6.

25. Rhie, A.; McCarthy, S.A.; Fedrigo, O.; Damas, J.; Formenti, G.; Koren, S.; Uliano-Silva, M.; Chow, W.; Fungtammasan, A.; Kim, J.; et al. Towards complete and error-free genome assemblies of all vertebrate species. Nature 2021, 592, 737–746, doi:10.1038/s41586-021-03451-0.

26. Shearin, A.L.; Ostrander, E.A. Leading the way: canine models of genomics and disease. Dis Model Mech 2010, 3, 27–34, doi:10.1242/dmm.004358.

27. Karlsson, E.K.; Lindblad-Toh, K. Leader of the pack: gene mapping in dogs and other model organisms. Nat Rev Genet 2008, 9, 713–725, doi:10.1038/nrg2382.

28. Tkaczyk-Wlizlo, A.; Kowal, K.; Slaska, B. Mitochondrial DNA alterations in the domestic dog (Canis lupus familiaris) and their association with development of diseases: A review. Mitochondrion 2022, 63, 72–84, doi:10.1016/j.mito.2022.02.001.

29. Slaska, B.; Grzybowska-Szatkowska, L.; Surdyka, M.; Nisztuk, S.; Rozanska, D.; Rozanski, P.; Smiech, A.; Orzelski, M. Mitochondrial D-loop mutations and polymorphisms are connected with canine malignant cancers. Mitochondrial DNA 2014, 25, 238–243, doi:10.3109/19401736.2013.792054.

30. Slaska, B.; Grzybowska-Szatkowska, L.; Nisztuk, S.; Surdyka, M.; Rozanska, D. Mitochondrial DNA polymorphism in genes encoding ND1, COI and CYTB in canine malignant cancers. Mitochondrial DNA 2015, 26, 452–458, doi:10.3109/19401736.2013.840594.

31. Strakova, A.; Ni Leathlobhair, M.; Wang, G.D.; Yin, T.T.; Airikkala-Otter, I.; Allen, J.L.; Allum, K.M.; Bansse-Issa, L.; Bisson, J.L.; Castillo Domracheva, A.; et al. Mitochondrial genetic diversity, selection and recombination in a canine transmissible cancer. Elife 2016, 5, doi:10.7554/eLife.14552.

32. Rebbeck, C.A.; Leroi, A.M.; Burt, A. Mitochondrial capture by a transmissible cancer. Science 2011, 331, 303, doi:10.1126/science.1197696.

33. Strakova, A.; Nicholls, T.J.; Baez-Ortega, A.; Ni Leathlobhair, M.; Sampson, A.T.; Hughes, K.; Bolton, I.A.G.; Gori, K.; Wang, J.; Airikkala-Otter, I.; et al. Recurrent horizontal transfer identifies mitochondrial positive selection in a transmissible cancer. Nat Commun 2020, 11, 3059, doi:10.1038/s41467-020-16765-w.

34. Ishiguro, N.; Nakajima, A.; Horiuchi, M.; Shinagawa, M. Multiple nuclear pseudogenes of mitochondrial DNA exist in the canine genome. Mamm Genome 2002, 13, 365–372, doi:10.1007/s00335-001-2139-2.

35. Hazkani-Covo, E.; Zeller, R.M.; Martin, W. Molecular poltergeists: mitochondrial DNA copies (numts) in sequenced nuclear genomes. PLoS Genet 2010, 6, e1000834, doi:10.1371/journal.pgen.1000834.

36. Verscheure, S.; Backeljau, T.; Desmyter, S. In silico discovery of a nearly complete mitochondrial genome Numt in the dog (Canis lupus familiaris) nuclear genome. Genetica 2015, 143, 453–458, doi:10.1007/s10709-015-9844-3.

37. Wang, C.; Wallerman, O.; Arendt, M.L.; Sundstrom, E.; Karlsson, A.; Nordin, J.; Makelainen, S.; Pielberg, G.R.; Hanson, J.; Ohlsson, A.; et al. A novel canine reference genome resolves genomic architecture and uncovers transcript complexity. Commun Biol 2021, 4, 185, doi:10.1038/s42003-021-01698-x.

38. Meadows, J.R.S.; Kidd, J.M.; Wang, G.D.; Parker, H.G.; Schall, P.Z.; Bianchi, M.; Christmas, M.J.; Bougiouri, K.; Buckley, R.M.; Hitte, C.; et al. Genome sequencing of 2000 canids by the Dog10K consortium advances the understanding of demography, genome function and architecture. Genome Biol 2023, 24, 187, doi:10.1186/s13059-023-03023-7.

39. Simone, D.; Calabrese, F.M.; Lang, M.; Gasparre, G.; Attimonelli, M. The reference human nuclear mitochondrial sequences compilation validated and implemented on the UCSC genome browser. BMC Genomics 2011, 12, 517, doi:10.1186/1471-2164-12-517.

40. Kim, K.S.; Lee, S.E.; Jeong, H.W.; Ha, J.H. The complete nucleotide sequence of the domestic dog (Canis familiaris) mitochondrial genome. Mol Phylogenet Evol 1998, 10, 210–220, doi:10.1006/mpev.1998.0513.

41. Camacho, C.; Coulouris, G.; Avagyan, V.; Ma, N.; Papadopoulos, J.; Bealer, K.; Madden, T.L. BLAST+: architecture and applications. BMC Bioinformatics 2009, 10, 421, doi:10.1186/1471-2105-10-421.

42. Schall, P.Z.; Winkler, P.A.; Petersen-Jones, S.M.; Yuzbasiyan-Gurkan, V.; Kidd, J.M. Genome-wide methylation patterns from canine nanopore assemblies. G3 (Bethesda) 2023, 13, doi:10.1093/g3journal/jkad203.

43. Lindblad-Toh, K.; Wade, C.M.; Mikkelsen, T.S.; Karlsson, E.K.; Jaffe, D.B.; Kamal, M.; Clamp, M.; Chang, J.L.; Kulbokas, E.J., 3rd; Zody, M.C.; et al. Genome sequence, comparative analysis and haplotype structure of the domestic dog. Nature 2005, 438, 803–819, doi:10.1038/nature04338.

44. Jagannathan, V.; Hitte, C.; Kidd, J.M.; Masterson, P.; Murphy, T.D.; Emery, S.; Davis, B.; Buckley, R.M.; Liu, Y.H.; Zhang, X.Q.; et al. Dog10K_Boxer_Tasha_1.0: A Long-Read Assembly of the Dog Reference Genome. Genes (Basel) 2021, 12, doi:10.3390/genes12060847.

45. Field, M.A.; Yadav, S.; Dudchenko, O.; Esvaran, M.; Rosen, B.D.; Skvortsova, K.; Edwards, R.J.; Keilwagen, J.; Cochran, B.J.; Manandhar, B.; et al. The Australian dingo is an early offshoot of modern breed dogs. Sci Adv 2022, 8, eabm5944, doi:10.1126/sciadv.abm5944.

46. Field, M.A.; Rosen, B.D.; Dudchenko, O.; Chan, E.K.F.; Minoche, A.E.; Edwards, R.J.; Barton, K.; Lyons, R.J.; Tuipulotu, D.E.; Hayes, V.M.; et al. Canfam_GSD: De novo chromosome-length genome assembly of the German Shepherd Dog (Canis lupus familiaris) using a combination of long reads, optical mapping, and Hi-C. Gigascience 2020, 9, doi:10.1093/gigascience/giaa027.

47. Halo, J.V.; Pendleton, A.L.; Shen, F.; Doucet, A.J.; Derrien, T.; Hitte, C.; Kirby, L.E.; Myers, B.; Sliwerska, E.; Emery, S.; et al. Long-read assembly of a Great Dane genome highlights the contribution of GC-rich sequence and mobile elements to canine genomes. Proc Natl Acad Sci U S A 2021, 118, doi:10.1073/pnas.2016274118.

48. Sinding, M.S.; Gopalakrishnan, S.; Raundrup, K.; Dalen, L.; Threlfall, J.; Darwin Tree of Life Barcoding, c.; Wellcome Sanger Institute Tree of Life, p.; Wellcome Sanger Institute Scientific Operations, D.N.A.P.c.; Tree of Life Core Informatics, c.; Darwin Tree of Life, C.; et al. The genome sequence of the grey wolf, Canis lupus Linnaeus 1758. Wellcome Open Res 2021, 6, 310, doi:10.12688/wellcomeopenres.17332.1.

49. Bredemeyer, K.R.; vonHoldt, B.M.; Foley, N.M.; Childers, I.R.; Brzeski, K.E.; Murphy, W.J. The value of hybrid genomes: Building two highly contiguous reference genome assemblies to advance Canis genomic studies. J Hered 2024, 115, 480–486, doi:10.1093/jhered/esae013.

50. Bjornerfeldt, S.; Webster, M.T.; Vila, C. Relaxation of selective constraint on dog mitochondrial DNA following domestication. Genome Res 2006, 16, 990–994, doi:10.1101/gr.5117706.

51. Li, H. Minimap2: pairwise alignment for nucleotide sequences. Bioinformatics 2018, 34, 3094–3100, doi:10.1093/bioinformatics/bty191.

52. Li, H. New strategies to improve minimap2 alignment accuracy. Bioinformatics 2021, 37, 4572–4574, doi:10.1093/bioinformatics/btab705.

53. Quinlan, A.R.; Hall, I.M. BEDTools: a flexible suite of utilities for comparing genomic features. Bioinformatics 2010, 26, 841–842, doi:10.1093/bioinformatics/btq033.

54. Peng, Y.; Li, H.; Liu, Z.; Zhang, C.; Li, K.; Gong, Y.; Geng, L.; Su, J.; Guan, X.; Liu, L.; et al. Chromosome-level genome assembly of the Arctic fox (Vulpes lagopus) using PacBio sequencing and Hi-C technology. Mol Ecol Resour 2021, 21, 2093–2108, doi:10.1111/1755-0998.13397.

55. Laidre, K.L.; Supple, M.A.; Born, E.W.; Regehr, E.V.; Wiig, O.; Ugarte, F.; Aars, J.; Dietz, R.; Sonne, C.; Hegelund, P.; et al. Glacial ice supports a distinct and undocumented polar bear subpopulation persisting in late 21st-century sea-ice conditions. Science 2022, 376, 1333–1338, doi:10.1126/science.abk2793.

56. Buckley, R.M.; Davis, B.W.; Brashear, W.A.; Farias, F.H.G.; Kuroki, K.; Graves, T.; Hillier, L.W.; Kremitzki, M.; Li, G.; Middleton, R.P.; et al. A new domestic cat genome assembly based on long sequence reads empowers feline genomic medicine and identifies a novel gene for dwarfism. PLoS Genet 2020, 16, e1008926, doi:10.1371/journal.pgen.1008926.

57. Kent, W.J. BLAT--the BLAST-like alignment tool. Genome Res 2002, 12, 656–664, doi:10.1101/gr.229202. Article published online before March 2002.

58. Tsuji, J.; Frith, M.C.; Tomii, K.; Horton, P. Mammalian NUMT insertion is non-random. Nucleic Acids Res 2012, 40, 9073–9088, doi:10.1093/nar/gks424.

59. Trumpff, C.; Michelson, J.; Lagranha, C.J.; Taleon, V.; Karan, K.R.; Sturm, G.; Lindqvist, D.; Fernstrom, J.; Moser, D.; Kaufman, B.A.; et al. Stress and circulating cell-free mitochondrial DNA: A systematic review of human studies, physiological considerations, and technical recommendations. Mitochondrion 2021, 59, 225–245, doi:10.1016/j.mito.2021.04.002.

60. Tao, Y.; He, C.; Lin, D.; Gu, Z.; Pu, W. Comprehensive Identification of Mitochondrial Pseudogenes (NUMTs) in the Human Telomere-to-Telomere Reference Genome. Genes (Basel) 2023, 14, doi:10.3390/genes14112092.

61. Lee, B.T.; Barber, G.P.; Benet-Pages, A.; Casper, J.; Clawson, H.; Diekhans, M.; Fischer, C.; Gonzalez, J.N.; Hinrichs, A.S.; Lee, C.M.; et al. The UCSC Genome Browser database: 2022 update. Nucleic Acids Res 2022, 50, D1115–D1122, doi:10.1093/nar/gkab959.

62. Raney, B.J.; Dreszer, T.R.; Barber, G.P.; Clawson, H.; Fujita, P.A.; Wang, T.; Nguyen, N.; Paten, B.; Zweig, A.S.; Karolchik, D.; et al. Track data hubs enable visualization of user-defined genome-wide annotations on the UCSC Genome Browser. Bioinformatics 2014, 30, 1003–1005, doi:10.1093/bioinformatics/btt637.

63. Nguyen, A.K.; Blacksmith, M.S.; Kidd, J.M. Duplications and Retrogenes Are Numerous and Widespread in Modern Canine Genomic Assemblies. Genome Biol Evol 2024, 16, doi:10.1093/gbe/evae142.

