## Supplementary Figures for "Characterization of nuclear mitochondrial insertions in canine genome assemblies"

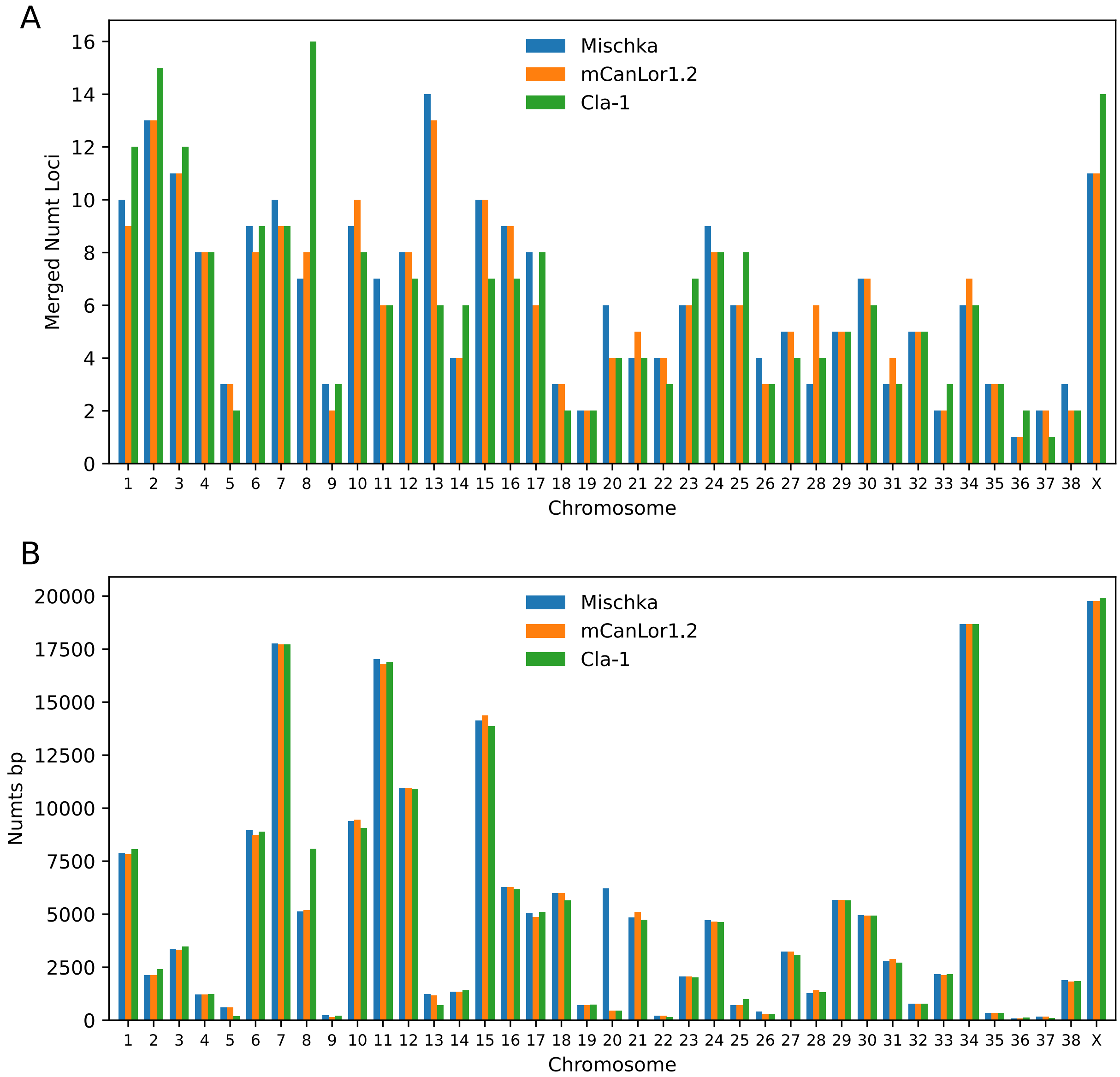


**Figure S1.** Numts identified per chromosome

A barplot of the total number of (A) the merged Numts loci and (B) total length of Numts segments identified per chromosome is shown for the Mischka (German Shepherd), mCanLor1.2 (Greenland wolf), and Cla-1 (coyote) assemblies. Numts length is calculated based on the size of the aligned HSP along the mitochondria genome.

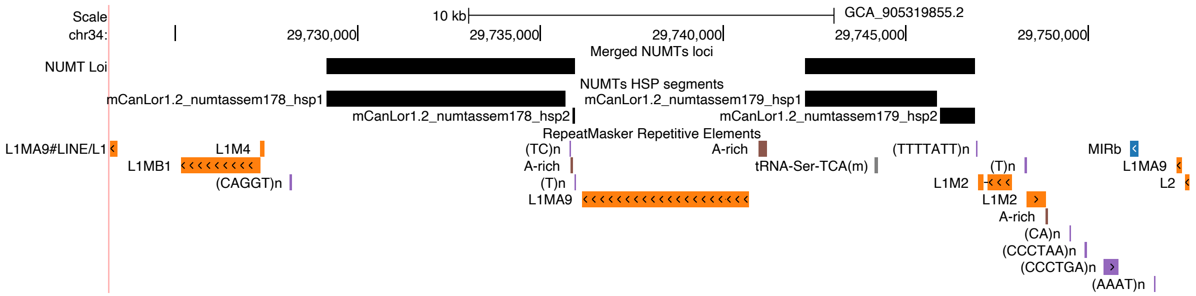


**Figure S2.** A long Numt is disrupted by a LINE-1 insertion in the mCanLor1.2 assembly
A UCSC genome browser view is shown for a region on mCanLor1.2 chr32, which corresponds to chr34:12617928-12629146 in the Mischka genome assembly. The two Numt HSP segments that are disrupted in this wolf assembly are present as a contiguous segment in Mischka.

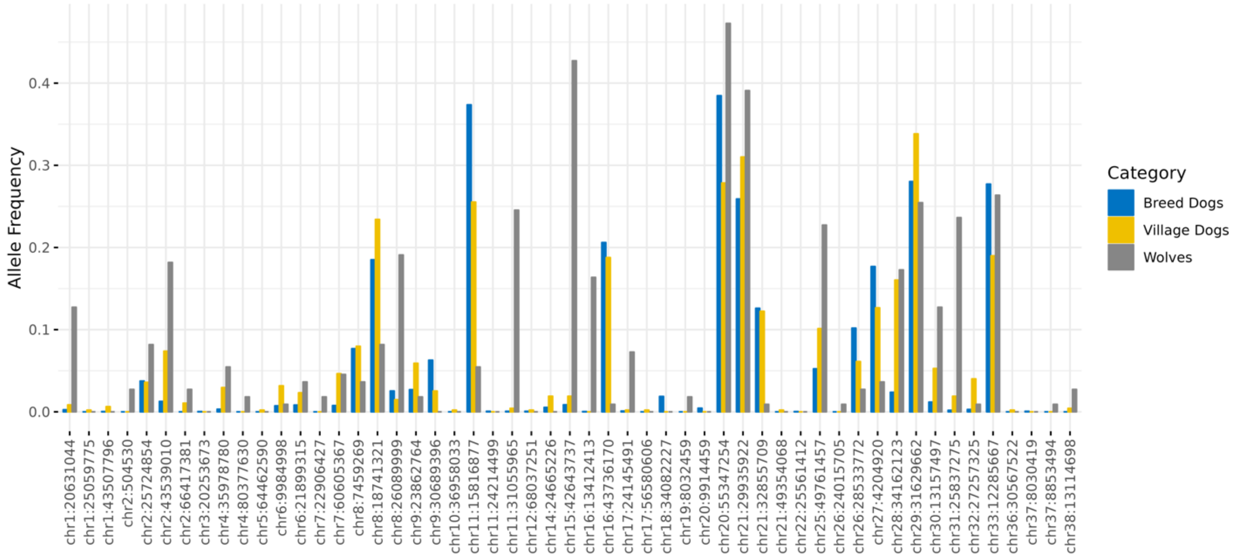


**Figure S3.** Allele frequency of 53 Numt insertions identified in Dog10K samples.
The frequency of each inserted is plotted for Breed Dogs (blue), Village Dogs (yellow) and Wolves (gray).
